# Both ANT and ATPase are essential for mitochondrial permeability transition but not depolarization

**DOI:** 10.1101/2021.11.16.468842

**Authors:** M.A. Neginskaya, S.E. Morris, E.V. Pavlov

## Abstract

A sudden increase in permeability of the mitochondrial inner membrane, mitochondrial permeability transition (PT), is the central event responsible for cell death and tissue damage in conditions such as stroke and heart attack. PT is caused by the opening of the Cyclosporin A (CSA) dependent calcium-induced pore, the Permeability Transition Pore (PTP). The molecular details of PTP are incompletely understood. We utilized a combination of holographic and fluorescent microscopy to assess the contribution of the ATP synthase and Adenine Nucleotide Translocator (ANT) towards PTP. In cells lacking either ATP synthase or ANT, we observed CSA-sensitive membrane depolarization, but not high-conductance PTP. Further, we found that in wild-type cells calcium induced CSA-sensitive depolarization precedes opening of the PTP, which occurred until after nearly complete mitochondrial membrane depolarization. We propose that both ATP synthase and ANT are required for high conductance PTP but not depolarization, which presumably occurs through activation of the low conductance PT, which has a molecular nature that is different from both complexes.

## Introduction

The low permeability of the mitochondrial inner membrane is an essential condition for efficient coupling between respiratory chain activity and phosphorylation of ADP by ATP synthase (Boyer, 1997; Mitchell, 1961). An increase in the permeability of the inner membrane leads to mitochondrial membrane depolarization, uncoupling of the oxidative phosphorylation, and mitochondrial energy failure. It is generally accepted that stress-induced increase in the permeability of the mitochondrial inner membrane, known as permeability transition (PT) is a critical contributor towards cell death in a wide range of pathologies associated with hypoxic-ischemic injuries (Bernardi et al., 2006). PT is caused by the activation of the PT pore (PTP) in the mitochondrial inner membrane.

The signature feature of PTP is an unselective increase in membrane permeability to ions and other molecules up to 1.5 kDa in size, which can be blocked by Cyclosporin A (CSA) (Crompton et al., 1988). The molecular mechanisms of PTP are not entirely understood and are the subject of intensive investigation and considerable controversies (Alavian et al., 2014; Bonora et al., 2013; Carroll et al., 2019; Giorgio et al., 2013; He et al., 2017b; Karch et al., 2019; Neginskaya et al., 2019). Genetic knock-out studies suggest that PTP involves both Adenine Nucleotide Translocator (ANT) and ATP synthase (ATPase) (Bonora et al., 2022; Bround et al., 2020). However, since both ANT (Brustovetsky and Klingenberg, 1996) and ATPase (Alavian *et al*., 2014; Giorgio *et al*., 2013) can form the pore when purified from the mitochondria and reconstituted into model membranes, the question about which of these channels is responsible for PTP formation remains open.

We reasoned that such a controversy could be explained by the lack of unambiguous methodology to measure PT inside the living cells. Despite the large arsenal of methods available to experimentally study PTP, the number of direct assays in the intact cells is surprisingly limited, with many conclusions regarding PTP activity derived from the measurements of the mitochondrial membrane depolarization (Carroll *et al*., 2019).

Since depolarization is not necessarily caused by the PTP this method often leads to inconclusive results and interpretations. To overcome this problem, we developed a novel assay that is based on the holographic imaging technique (Sandoz et al., 2019). This assay allows direct detection of mitochondrial membrane permeabilization (and hence PTP) inside the living cells. A holographic microscope generates images based on the differences in refractive indexes (RI) of the object parts. RI reflects how fast the light propagates through the object (mitochondrion in the case of this study). By estimation of the delay of the light passing through the matrix of the intact mitochondria with higher RI the holographic microscope can reconstruct their shape (Fig.1). Due to the large size of PTP the immediate consequence of its opening is a rapid exchange of the solute contents across the mitochondrial inner membrane. This exchange causes equilibration of the solute content and results in the equalization of refractive indexes (RI) of the mitochondrial matrix and its surroundings (Fig. 1C) and as a result, disappearance of the organelles from holographic image (RI image).

**Figure 1.**
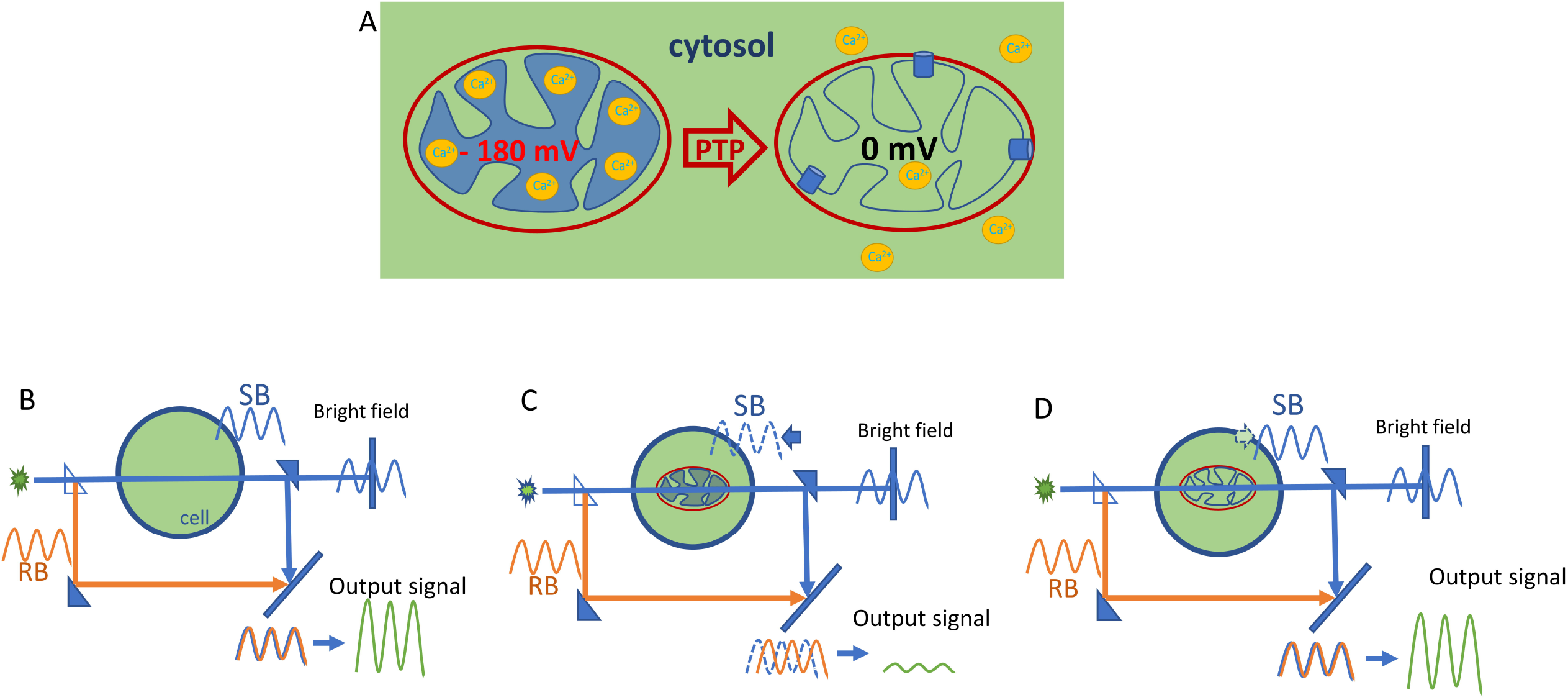
Principles of mitochondrial permeability transition assay in living cells using holographic microscopy. *A. PTP causes the equalization of RIs between mitochondrial matrix and cytosol*. Mitochondrial Ca^2+^ overload leads to the induction of the mitochondrial membrane permeabilization that occurs due to the opening of the PTP in the inner mitochondrial membrane. PTP opening causes equilibration of solutes of up to 1.5 kDa in size between matrix and cytosol. Equilibration of solutes results in equalization of RIs (and ODs) between mitochondrial matrix and cytosol. *B. Principle of the holographic imaging*. A conventional bright-field microscope detects only the signal that is passed through the sample, sample beam (SB). Holographic imaging detects the interference pattern of 2 light beams: the reference beam (RB) and SB, which passes through the object of interest. SB is initially equal to RB and becomes delayed by passing through the sample. This delay of SB depends on the RIs of the content inside the sample. In the present example, SB and RB reach the detector in the same phase, and the holographic microscope detector registers the interference, which will be the sum of two beams with equal phase and amplitude. C. Intact mitochondrion has different RI (higher optical density, OD) when compared to the cytosol of the cell. If cellular cytosol contains intact mitochondrion with higher OD, the SB is delayed due to the slower light speed inside the matrix compared to the RB. As a result, the interference pattern will be changed because of the differences between the phases of RB and SB. A Bright-field microscope is not able to distinguish the SB with or without mitochondrion on its way as mitochondrion is transparent and does not cause enough of decrease in intensity of the light. D. Following the PTP opening RI (and OD) of the mitochondrial matrix is equalized with RI of the cytosol of the cell and does not cause the delay of SB. In this case, the interference pattern detected by the microscope will be the same to that of the cytosol without mitochondrion (compare outcome signal in panel B). As a result, mitochondrion with PTP opened becomes “invisible” for the holographic microscope.

Here, we use a combination of fluorescent and holographic microscopy to simultaneously measure the changes of mitochondrial membrane potential and PTP activation in wild-type cells, as well as in cells lacking either ATPase or ANT.

We discovered that calcium-induced high-conductance PTP is preceded by the initial stage of membrane depolarization (which we define as low-conductance Permeability Transition). Further, we demonstrate that, deletion of either ATPase or ANT leads to complete elimination of PTP. Interestingly, neither of these proteins was essential for CSA sensitive calcium-induced loss of membrane potential. We hypothesize that activation of PTP requires cooperative molecular interactions of ANT and ATPase.

## Materials and methods

### Cell culture

Immortalized HAP 1 and MEF cell lines were used for holographic imaging assay. The cells were cultured as described previously (He *et al*., 2017b; Karch *et al*., 2019). Briefly, HAP 1 cells were grown Iscove’s Modified Dulbecco’s Medium (IMDM), supplemented with 10% Heat-Inactivated Fetal Bovine Serum (HI FBS; Life Technologies), 10 mL per L of Antibiotic Antimycotic Solution (Penicillin/Streptomycin/Amphoterichin B; Sigma Aldrich) and 2 mM L-Glutamine. MEF cells were grown in high-glucose Dulbecco’s Modified Eagle Medium (DMEM; Cytiva) supplemented with 10% Heat-Inactivated Fetal Bovine Serum (HI FBS; Life Technologies), 10 mL per L of Antibiotic Antimycotic Solution (Penicillin/Streptomycin/Amphoterichin B; Sigma Aldrich), 1X Non-Essential Amino acids (NEAA; Lonza). Cells were maintained in a humidified cell incubator, at 37°C under a 5% CO2 atmosphere. HAP 1 Δ (c+δ) cell line that lacks c and δ-subunits of ATP synthase were used for the study of the role of ATP synthase in high-conductance PTP. MEF ANT Triple KO cell line lacking 3 ANT genes was used to study the role of ANT in high conductance PTP. MEF ANT Triple KO cells were grown in the same media as MEF WT cells with the addition of 1mM Sodium Pyruvate (Gibco) and 25mg/500mL Uridine (Sigma Aldrich).

### Holographic and Fluorescent Imaging

The cells were plated on poly-D-lysine coated glass coverslips 24 hours before imaging to reach the confluency of 70-90 %. Before the experiment, the coverslips with the cells were placed in the imaging chamber and washed with Hank’s Balanced Salt Solution (HBSS, Gibco). TMRM fluorescent probe (Invitrogen) was used for estimation of mitochondrial membrane potential. Cells were incubated with 40 nM of TMRM for 15 min in room temperature in the darkness. Recording media contained 40 nM of TMRM. Ferutinin (Sigma Aldrich) was used to induce calcium-induced PT. Holographic (RI) images and TMRM signal were acquired every 15 seconds with aid of 3D Cell Explorer-fluo (Nanolive, Switzerland) equipped with 60X objective. Protonophore FCCP (10 µM; Sigma Aldrich) was used at the end to observe the drop of membrane potential and normalize the TMRM signal.

### Data processing and analysis

Fiji ImageJ was used to process holographic reconstructions. Multipage TIF files were prepared as described in (Cotte et al., 2013). Plain RI images were reconstructed as a Z-stack maximal intensity projection from the volume of the cell that contained mitochondria. Ilastik, the interactive learning and segmentation toolkit, was used for mitochondrion segmentation. After being trained by the user, Ilastik tool creates the probability map of pixels that relate to mitochondria and based on the probability, classify them as mitochondria (Fig. S1). Segmented images were converted to a binary image with Fiji “Make binary” tool. Resulted image is shown in Figure S2C.

To analyze the mitochondrial membrane permeabilization, we estimated the decrease in the refractive index (RI) of mitochondria by the decrease of the area occupied by mitochondria in reconstructed images. To do so, the regions of interest (ROIs) with functional mitochondria with maintained membrane potential were selected manually from the corresponding fluorescent images of cells labeled with TMRM (Fig. S1D).

These ROIs were used to estimate changes in membrane potential and applied to binary segmented masks created as described above (Fig. S1C). Next, the area occupied by mitochondria was estimated in selected ROIs in each time frame. The decrease of the area indicated the decrease of mitochondrial RI and, thus, mitochondrial permeabilization (Fig. S2).

Single mitochondrion tracking was performed manually by selecting the mitochondrion areas in RI images frame by frame. The same areas were used in corresponding TMRM fluorescent images to track the changes in mitochondrial membrane potential.

### Seahorse assay

Analysis of mitochondrial functions in HAP 1 WT and HAP 1 Δ(c+δ) cells was performed on Seahorse XFe24 (Agilent Technologies, USA) as described previously (Nichols et al., 2017). Briefly, cells were plated on Seahorse XFe24 Cell culture 24-well microplates 24 hours before experiment to reach the confluency 70-80% according to Agilent Technologies recommendations.

The night before the experiment, the cartridge containing the sensors was hydrated with 1 ml of XF Calibrant Solution per well and kept overnight in a CO_2_-free incubator. The day of the experiment, cells were washed with Seahorse XF DMEM medium that contained 1 mM pyruvate, 2 mM glutamine, and 10 mM glucose and incubated for 1 h in the CO_2_-free incubator (hypoxia). The cartridge was loaded with 30 µM of Ferutinin, 1 µM of FCCP and 0.5 µM of rotenone/antimycin A (Rot/AA) to the ports A, B and C accordingly to measure OCRs and ECARs. All the drugs were dissolved to the working concentrations using the Seahorse XF DMEM medium. Subsequently, the cells were loaded onto the analyzer and the measurements were conducted. The obtained data was exported and analyzed using the Seahorse Wave Desktop Software, which was downloaded from the Agilent Technologies’ website.

### Statistics

Origin 2021b software (OriginLab, Massachusetts, USA) was used for data presentation, analysis and statistics. All the data presented as Mean±SEM. The exact numbers of experiments (N) and cells (n) analyzed are mentioned in corresponding parts of the text. ANOVA and t-test were used to verify statistical significance.

## Results

### Visualization of mitochondria inside the living cells

In a holographic image, the contrast is achieved based on the differences in the refractive indexes (RI) of different areas of the cell (Cotte *et al*., 2013). Figure 2 illustrates that holographic imaging allows for the distinguishing of inner cellular structures that are not visible when bright-field imaging is applied (compare images on Figs. 2A and 2B). As shown in the RI image in Figures 2B and 2F, mitochondria (arrows) are visible directly inside the living cell without the use of fluorescent labels.

**Figure 2.**
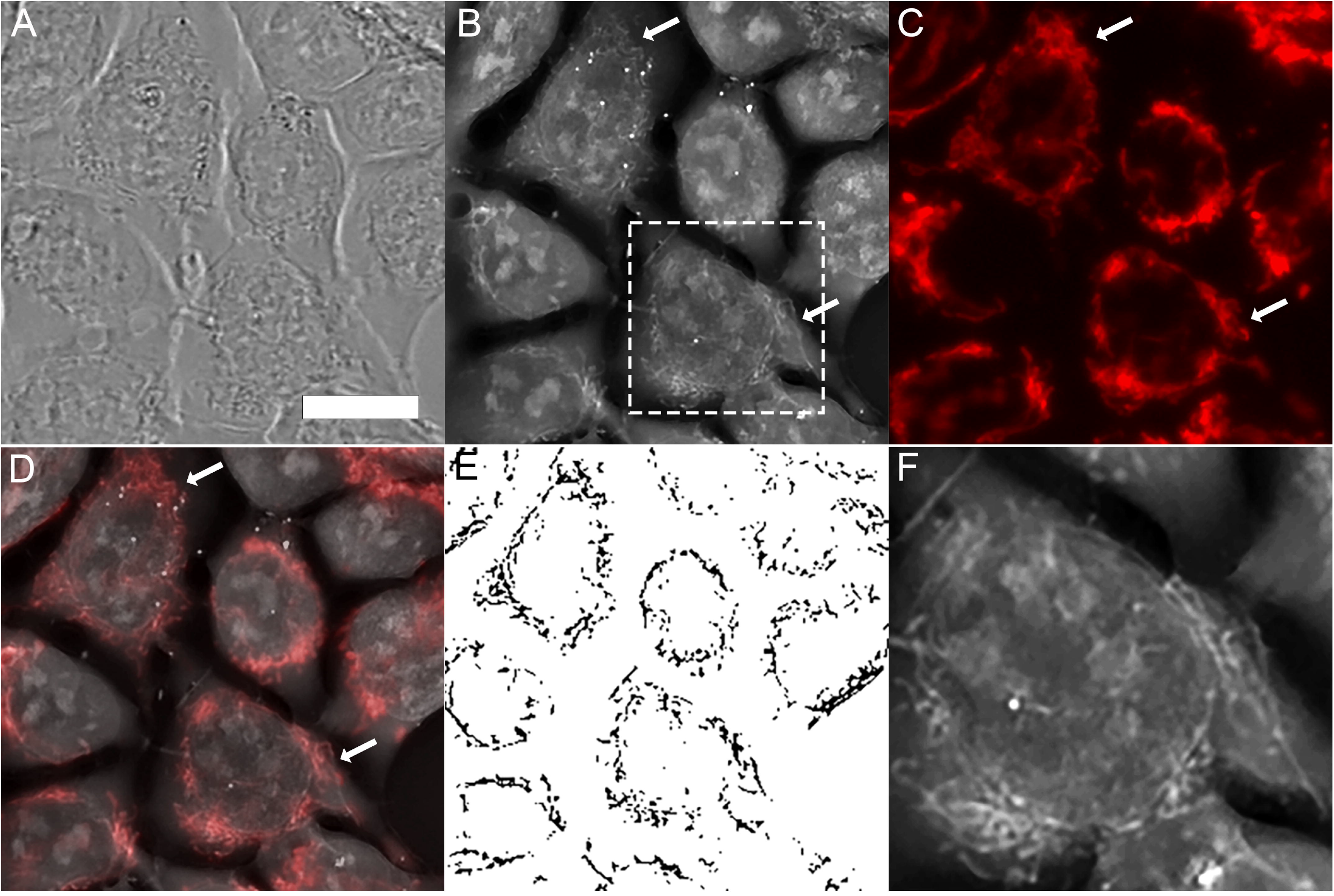
Holographic imaging allows monitoring of mitochondria in the living cells label-free. A. Bright field image of immortalized HAP-1 cell culture. B. Holographic image of the same area of the cells. No label was used. C. Staining of mitochondria with membrane potential sensitive probe TMRM. Areas with live mitochondria that maintain membrane potential are visible in fluorescent microscope. Arrows point to mitochondrial areas. D, Overlay, of the images shown on panels B and C showing colocalization between RI (B) and TMRM images (C). E. Segmentation of the mitochondrial regions from the holographic image shown on panel B. These segmented images were used for quantification of the PTP activation. F. Enlarged images of the region shown in the white box on panel B. Scale bar: 10 µm for A-E; 5 µm for F.

The identity of these structures was confirmed by the fluorescent probe TMRM, which selectively labels polarized mitochondria (Fig. 2C). Overlay of the RI and TMRM images allowed us to clearly identify structures representing mitochondria (Fig. 2 D). Using segmentation tool, we convert holographic images to binary mitochondrial maps (Fig. 2E). These maps were used to track permeabilization of mitochondria following treatments.

### Ferutinin models mitochondrial PT

We used calcium ionophore Ferutinin to model PT conditions. Ferutinin was shown to electrogenically deliver calcium into mitochondria and induce calcium overload followed by CsA sensitive mitochondrial depolarization (Abramov and Duchen, 2003), representing a robust cell culture model for the investigation of the molecular details of PTP (Carroll *et al*., 2019; He et al., 2017a; He *et al*., 2017b; Walker et al., 2020).

We measured the response of mitochondria to the addition of ferutinin in HAP1 cells by simultaneously monitoring of the membrane potential and RI of the mitochondria (Fig. 3 A-E and Fig. 4F). As can be seen from the figures, activation of PTP with ferutinin (20 µM) leads to mitochondrial depolarization (Fig. 3C and 3D) and disappearance of the mitochondrial structures from the RI images (Fig. 3A and 3B), which is consistent with the equilibration of the solutes and thus optical densities (and RI) between the matrix and cytoplasm. By segmenting mitochondria from other cellular structures and conversion of RI images to binary images (Fig. S2) we were able to track the drop in RI/”disappearance” of mitochondria with PTP as a decrease of the mitochondrial area on binary images (Fig. 3 E, black trace; Fig. S2). Drop in RI of mitochondria coincided with membrane depolarization that was detected by the decrease in TMRM signal (Fig. 3 E, Fig. 4F; N = 5; n = 82). Both membrane depolarization and RI drop were prevented by the addition of CSA (Fig. 3 F-J; Fig. 4F; N=4; n=89). These data demonstrate that non-selective mitochondrial membrane permeabilization can be directly detected in the living cells and that this increase in permeability is indicative of the activation of the CSA sensitive high-conductance PTP. In control experiments, when depolarization was induced by the protonophore FCCP without PTP induction, despite the drop in TMRM fluorescence mitochondria remained intact and easily recognizable in the RI images (Fig. 3 K-O, Fig. S3; N=3; n=55;) demonstrating that the detection of permeabilization does not depend on membrane potential.

**Figure 3.**
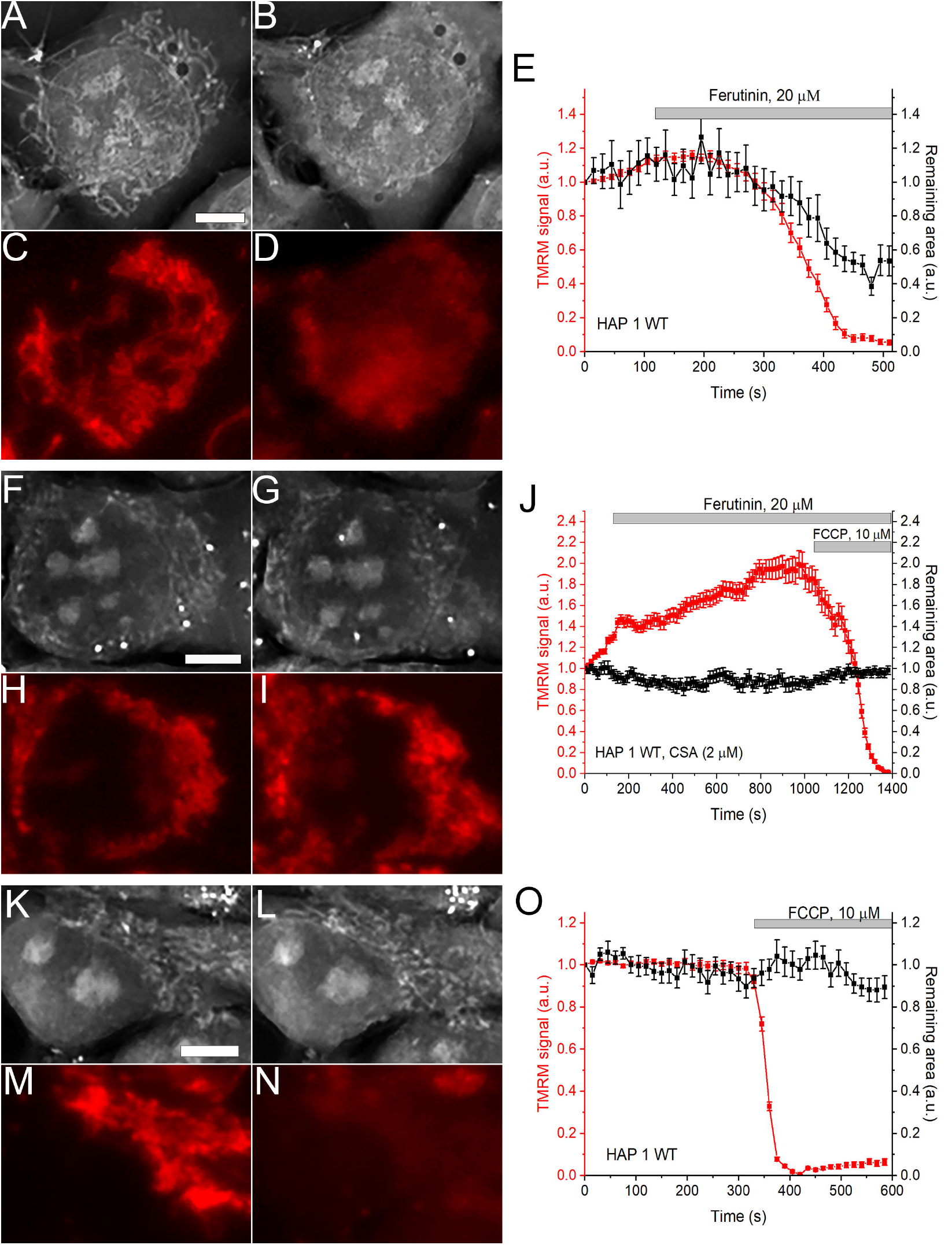
Simultaneous detection and quantification of the mitochondrial depolarization (TMRM, red) and permeabilization (RI) in the HAP 1 WT living cells. Panels A and B show holographic images of the cells before and 6 minutes after the addition of ferutinin (20 µM). Note the disappearance of the mitochondria at panel B. Panels C and D show fluorescent imaging of the mitochondrial membrane potential from the same field as in images A and B. Panels F-I, the same as in panels A-D but in the presence of CSA. Panels K-N – the same as in panels A-D but with addition of FCCP only. E, J, O time-dependent changes of TMRM and RI signals of the mitochondrial regions. Representative of experiment of N=5 for Ferutinin, N=4 for Ferutinin+CSA, N=3 for FCCP. Each data point represents value collected from different mitochondrial regions within the same field of view. Scale bar – 5 µm.

**Figure 4.**
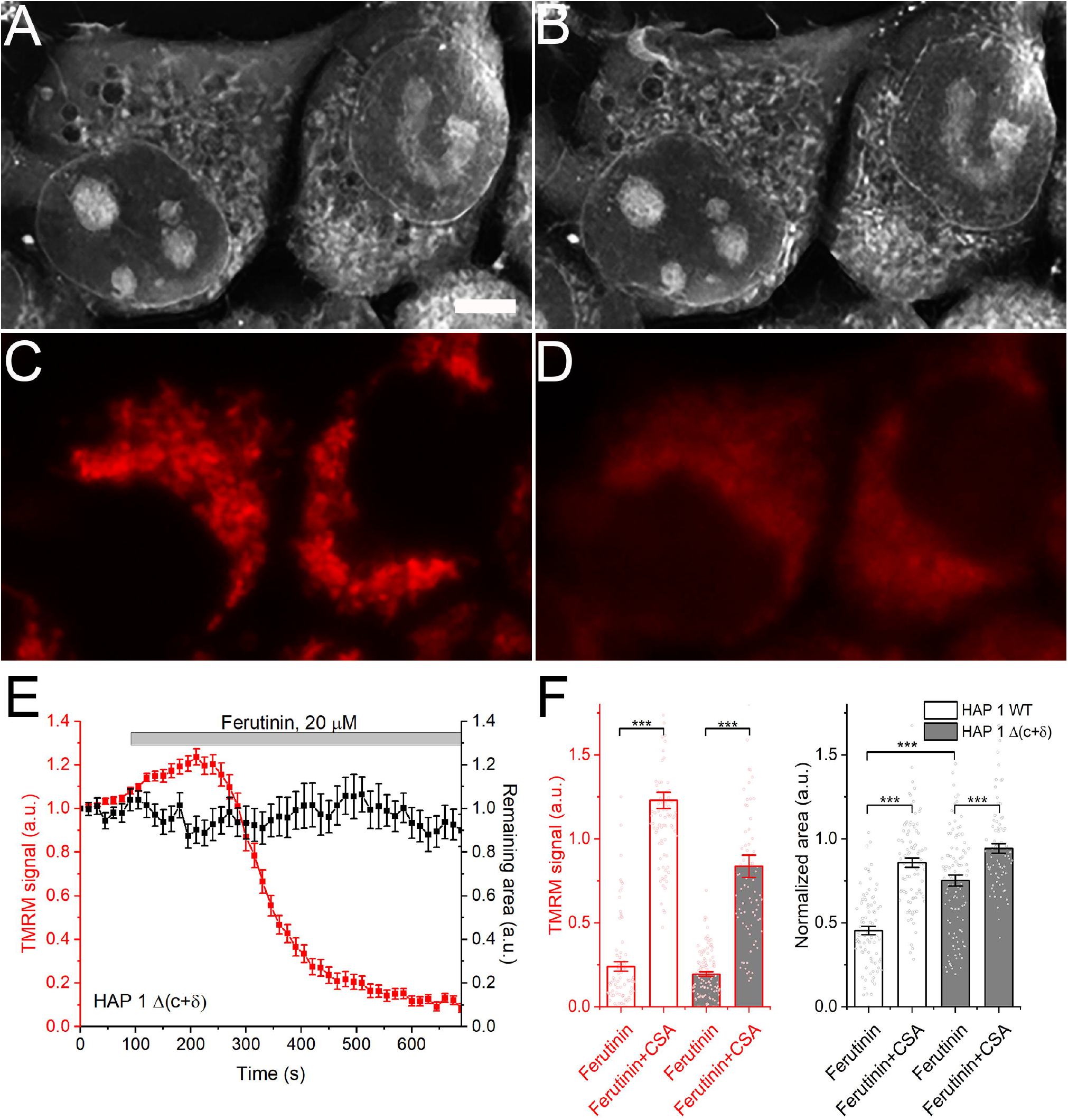
Lack of high-conductance PTP despite membrane depolarization in HAP1 Δ (c+δ) cells. Holographic and florescent (TMRM) images of cells before (A, C) and after (B, D) the addition of ferutinin (20 µM). Scale bar – 5 µm. E. Time dependence of the membrane depolarization and refractive index measurements. F. Quantification of the degrees of membrane depolarization and permeabilization. Mean±SEM; One way ANOVA; ***p<0.001

### Cells lacking assembled ATP synthase undergo CSA-sensitive depolarization but not membrane permeabilization

It has been suggested that ATP synthase plays an important role in the PTP. However, recent studies using a double knockout HAP1 mutant (HAP1 Δ (c+δ)), lacking c and δ subunits and, consequently, making them devoid of the assembled ATP synthase, show that these mitochondria can still undergo calcium-induced depolarization in the intact cells when stimulated with ferutinin (Carroll *et al*., 2019). Using holographic imaging, we investigated the relationship between the calcium-induced depolarization and high-amplitude mitochondrial permeabilization. As demonstrated in Figure 4, unlike wild-type cells, HAP1 Δ (c+δ) cells did not undergo high amplitude permeabilization, despite membrane depolarization (Fig. 4; N=5, n=108). This suggests that the assembled ATP synthase is required for the development of the high-conductance PTP but is not involved in the initial membrane depolarization that is triggered by the addition of ferutinin. Importantly, this initial depolarization was inhibited by CSA in both WT and HAP1 Δ (c+δ) (Fig. 4F), suggesting that it represents one of the stages of the PTP activation process.

The lack of high amplitude permeabilization was further confirmed by measurements of the effects of the ferutinin on mitochondrial respiration using Seahorse metabolic flux analyzer. As can be seen from Fig. 5 A, ferutinin caused rapid loss of mitochondrial function in the WT cells consistent with what would be expected from the high-conductance PTP activation and loss of the respiratory chain substrates. On the contrary, the same amount of ferutinin transiently stimulated mitochondrial respiration in the HAP1 Δ (c+δ) cells (Fig. 5 B). This stimulation of the respiration is consistent with the observation that despite depolarization, mitochondrial of these mutant cells remained structurally intact which allowed them to (at least transiently) maintain respiratory activity. The effects of the addition of ferutinin on the respiratory function for both cell types were blocked by CSA (Fig. 5), confirming that both processes, membrane depolarization and mitochondrial permeabilization, are related to PTP.

**Figure 5.**
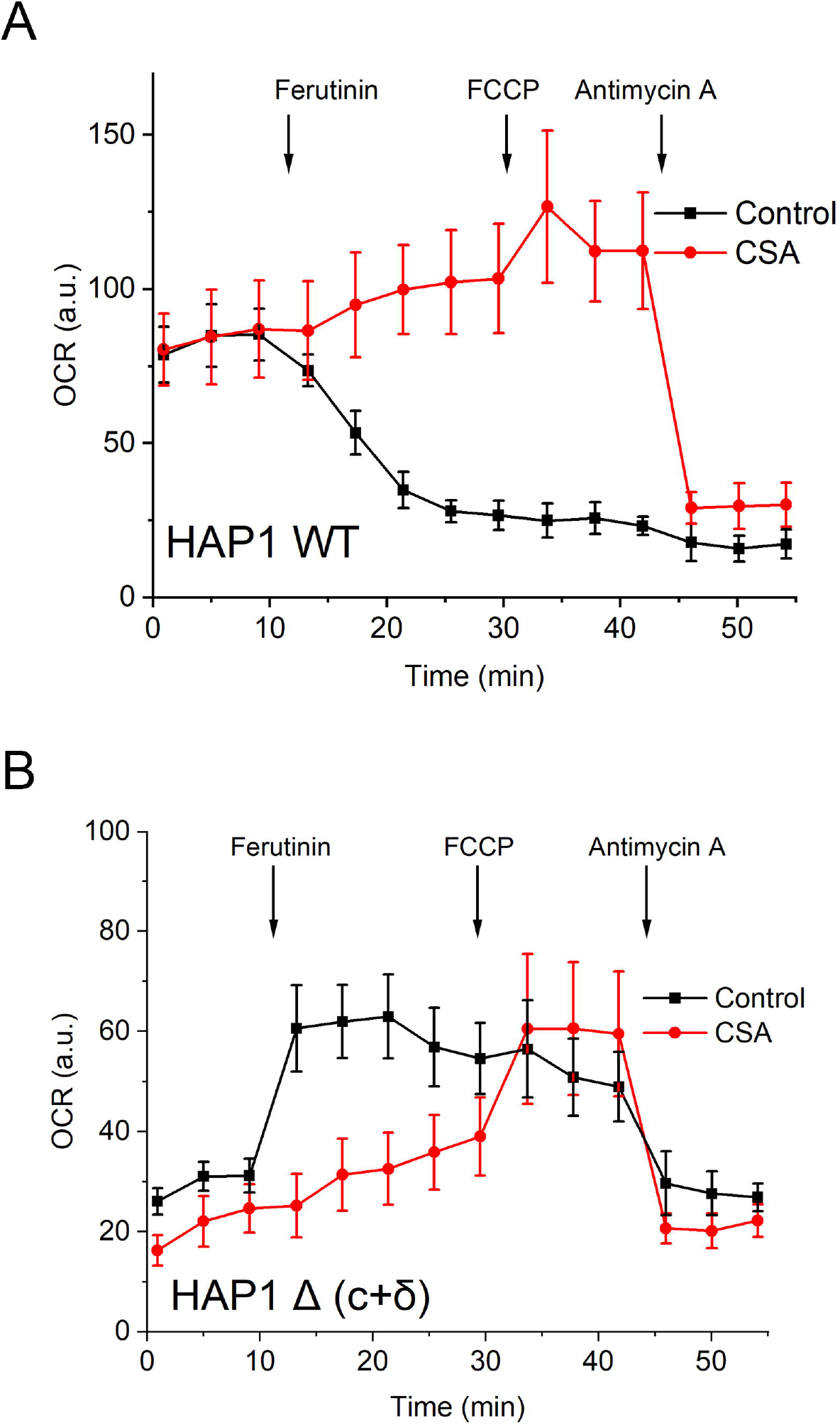
Seahorse analysis of the Oxygen consumption rates changes in response to ferutinin (30 µM). A. WT HAP 1 cells demonstrated dramatic decrease in respiratory rates that was prevented by the CSA (2 µM) consistent with the loss of mitochondrial function due to the PTP; B. HAP1 Δ (c+δ) cells respiration was stimulated by ferutinin (30 µM) but was not further stimulated by the uncoupler FCCP, consistent with the stimulation of the mitochondrial respiration due to Ca2+-induced uncoupling without high-amplitude PTP opening. This stimulation was prevented by the CSA (2 µM). Data are representative of 3 independent experiments.

### Cells lacking ANT undergo CSA-sensitive depolarization but not membrane permeabilization

Next, using holographic assay, we checked the permeabilization of mitochondria inside the MEF WT and ANT triple KO MEF cells upon ferutinin addition (30 µM for WT and 10 µM for ANT KO cells). Previous experiments showed that ANT triple KO MEF cells have significantly inhibited PTP (Karch *et al*., 2019). In WT MEF cells, decrease of the area occupied by mitochondria (Fig. 6 A,B,E) followed the depolarization induced by ferutinin (30 µM) addition (Fig.6 C,D,E; N=4; n=41). Like in case of HAP 1 cells, this process was inhibited by CSA, suggesting the involvement of PT (Fig. S4). However, in MEF ANT KO cells we did not observe any significant reduction in the area occupied by mitochondria followed by ferutinin (10 µM) treatment, while we still observed a dramatic loss of membrane potential (Fig. 6 F-J; N=5; n=56). Comparison of remaining mitochondrial area after ferutinin addition in WT and ANT KO cells is shown on Figure 6 K. These results suggest that like ATP synthase, ANT is also essential for the development of high-conductance PTP but not involved in calcium induced CSA sensitive mitochondrial depolarization.

**Figure 6.**
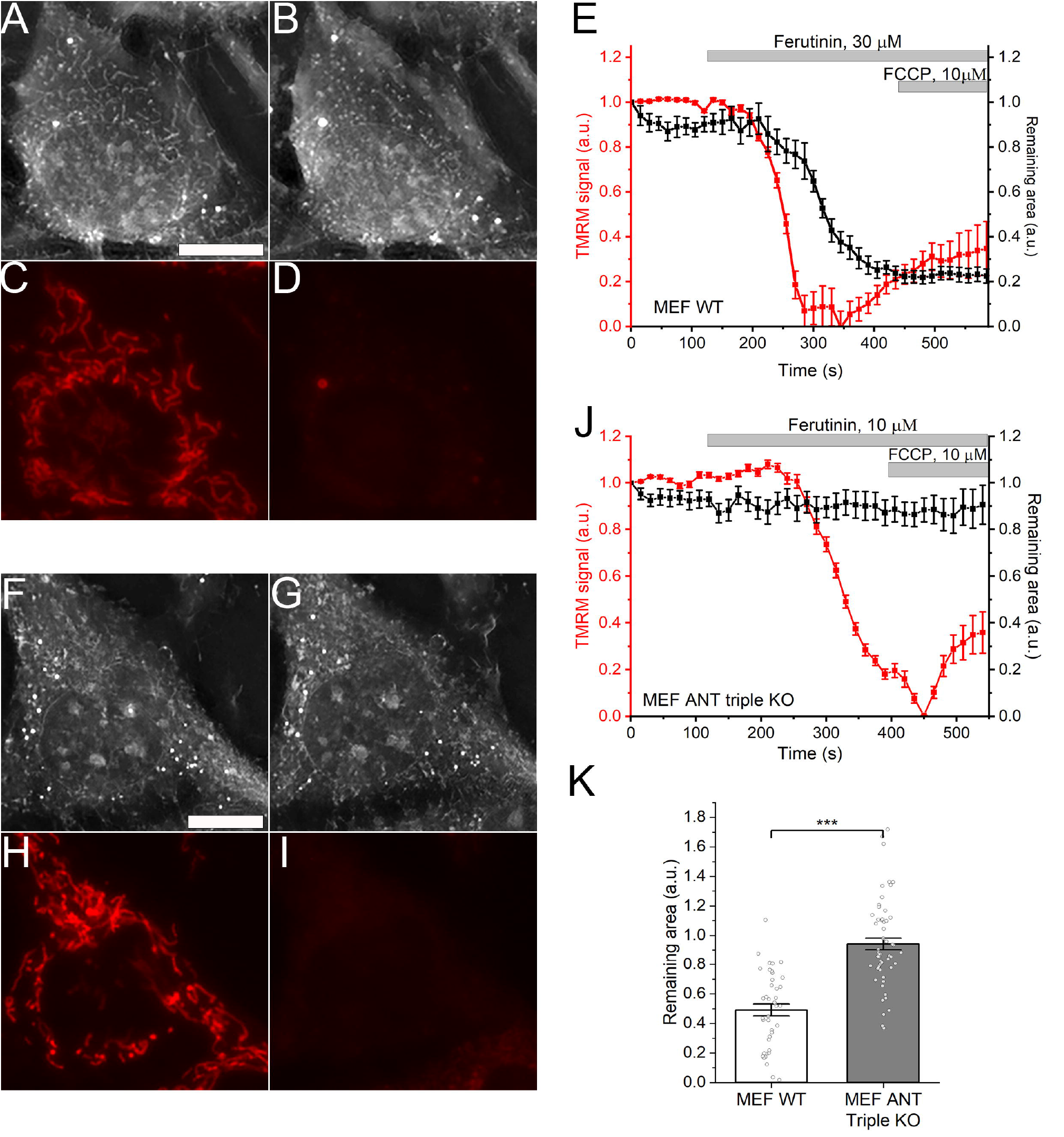
Lack of high-conductance PTP in ANT knock-out MEF cells. A-D simultaneous imaging of the RI and TMRM fluorescence before (A,C) and after addition of ferutinin (B,D) to the wild-type MEF cells. F-I, RI and TMRM fluorescence before (F,H) and after addition of ferutinin (G,I) to ANT triple KO MEF cells. Note that despite mitochondrial depolarization, mitochondria are still visible on RI images. E and J, time resolved quantification of the RI and TMRM signals following the addition of ferutinin. K, statistical analysis of the remaining mitochondrial area in RI images following ferutinin-induced depolarization. Note that mitochondria did not disappear in ANT triple KO MEF cells. N = 4 and n = 41 for WT; N = 5 and n = 56 for KO; Mean±SEM; One way ANOVA; ***p<0.001

### Mitochondrial calcium overload induces mitochondrial depolarization that precedes high-conductance permeabilization

In both mutant cell lines despite the lack of high-conductance PTP we observed mitochondrial depolarization. To gain further insight into the relationship between depolarization and permeabilization we analysed the dynamics of these two processes at the single mitochondrion level. Figure 7 shows the result of simultaneous analysis of the dynamics of mitochondrial membrane potential and non-selective permeabilization performed at the level of a single mitochondrion. Here, we traced individual mitochondria using both RI and TMRM readouts from the moment before treatment where mitochondria were functional (Fig. 7A) and visible on RI image (Fig. 7B) until the mitochondrial disappearance (Fig. 7D). The specific organelles RI were tracked throughout the duration of the experiment frame by frame as shown in Fig. 7C for 2 selected mitochondria. TMRM signal was detected at corresponding areas of fluorescent images (Fig. 7A). As shown in Figures 7E and 7H, ferutinin caused a gradual decrease in the membrane potential. Interestingly, despite significant membrane depolarization, the RI of individual mitochondrion stayed largely undisturbed and individual mitochondrion remained clearly visible (Figs. 7E and 7F for mitochondrion 1, and 7H and 7I for mitochondrion 2). However, mitochondria rapidly disappeared from RI images when depolarization was nearly complete (Figs 6E and 6G for mitochondrion 1, and 6H and 6J for mitochondrion 2). The membrane potential of individual mitochondrion at the moment of organelle disappearance from the RI image was 15 ± 6 % of the initial potential level (Fig. 7K, left panel, n=10). The average time delay from the start of depolarization until the disappearance of the individual mitochondrion was 150 ± 20 seconds (Fig. 7K, right panel, n=10; p<0.001). Overall, individual mitochondrion analysis showed that almost complete depolarization occurred prior to the onset of non-selective large-scale membrane permeabilization, and on average, the permeabilization was delayed by 150 ± 20 seconds from the beginning of the depolarization (Fig. 7K, n=10; p<0.001). Altogether, these experiments indicated that initial depolarization occurred prior to high-conductance PTP activation. This is a new insight that suggests that the high-conductance PTP is not the cause of membrane depolarization.

**Figure 7.**
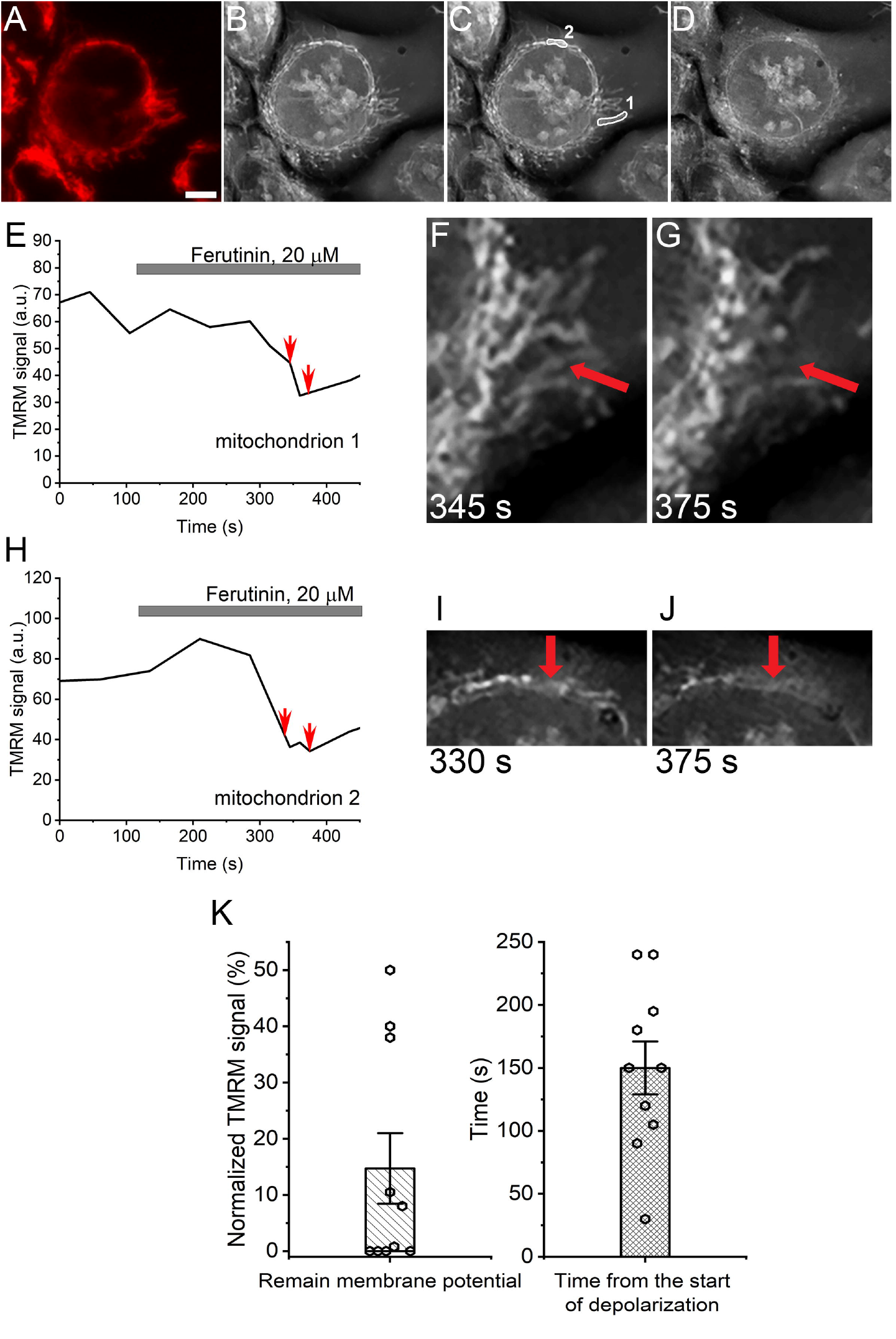
Monitoring time-dependent membrane depolarization and high-amplitude permeabilization in HAP 1 WT cells at the level of a single mitochondrion. A-D. Fluorescent (TMRM) (A) and holographic images of the cell at the beginning of the experiment (B,C) and following ferutinin addition (D). Labels on panel C show the selection of two representative mitochondria. Scale bar – 5 µm. E, H. time dependence of the TMRM fluorescence from the mitochondria #1 and #2 (see panel C). F, G images correspond to the time points marked by arrows at panel E for mitochondrion #1; note the disappearance of mitochondrion from the panel G. H-J analysis similar to the panels E-G for the mitochondrion #2. K. The relationship between mitochondrial depolarization and permeabilization at the level of single mitochondrion. Left panel, the level of the residual membrane potential at the moment of mitochondrial permeabilization (n=10). Right panel, the time delay between the offset of depolarization and permeabilization (n=10; p<0.001; t-test for null-hypothesis).

## Discussion

Traditionally, a functional assay of the PTP in intact cells relies on the fluorescent measurements of the mitochondrial parameters. In most cases, PTP can be experimentally identified as a calcium-induced CSA-sensitive membrane depolarization and/or calcium release, both of which can be detected fluorometrically in the intact cells (Bonora et al., 2016; Duchen, 2004). Notably, these methods do not necessarily indicate activation of the high-conductance PTP. To our knowledge, the only fluorescent method specifically geared towards PTP is monitoring the calcein release from the mitochondria (Petronilli et al., 1999), where calcein release would indicate the opening of the large pore. However, calcium triggered calcein (which is similar in size to the essential mitochondrial energy metabolite NADH known to be released through PTP) release can occur in a CSA independent manner and without the loss of mitochondrial function (Petronilli *et al*., 1999), suggesting that in addition to PTP this release can proceed through the mechanisms independent of the simple size-exclusion diffusion through the large pore. The method described here provides a direct assay that relies on the definitive feature of PTP, which does not rely on tracking of the transport of the specific molecule but rather reflects the non-selective equilibration of the solutes across the mitochondrial membrane.

Further, unlike in experiments involving isolated mitochondria in the population, we were able to monitor optical density (RI) in a single mitochondrion. This is an important advantage as changes in light scattering in the population of mitochondria might not necessarily reflect complete swelling of individual organelles, but rather gradual changes in “average” light scattering across the whole population. We anticipate that this method will help to clarify many details in the PT at the level of intact cells and resolve some current controversies.

One important aspect of PTP which this new method would allow for the clarification of, is the ability to more accurately estimate the relationship between PTP and mitochondrial swelling. It is known that, following calcium treatment, isolated mitochondria swell (Haworth and Hunter, 1979; Hunter et al., 1976) and, in the literature, generally the terms “light-scattering” and “swelling” assay are used interchangeably. However, prior to PTP opening, mitochondria are perfectly osmotically and oncotically balanced with the surrounding medium. Opening of the non-selective PTP - which allows flux of molecules of up to 1.5 kDa in size - would definitely cause a drop in RI due to the equilibration of the matrix and medium content. At the same time, however, this solute exchange should not necessarily lead to swelling in the living cell. The oncotic pressure of non-permeable proteins would remain balanced, as it was prior to PTP, while permeable molecules would exchange freely, leaving the net accompanying water flux unchanged. Single mitochondria RI imaging will help to clarify if swelling is indeed the direct consequence of the PTP opening, or if swelling occurs at the later stages of mitochondrion demise.

The advantage of being able to monitor RI in real-time with single organelle resolution is evident from our experiments with simultaneous monitoring of the RI in relation to the mitochondrial membrane potential. As shown in Figure 6, during the induction of the PTP by the addition of calcium, we detected that at the first stage mitochondria undergo membrane depolarization, followed by a second stage of the PTP characterized by high amplitude membrane permeabilization. This observation challenges the widely accepted view that calcium-induced PTP is a cause of membrane depolarization (Zoratti and Szabo, 1995). Our data suggest that the initial step of PTP activation is likely the opening of the lower conductance channel that is sufficient to depolarize mitochondria. This occurs prior to the activation of the high-conductance PTP which is required for mitochondrial swelling (as seen in the isolated mitochondria).

The molecular mechanisms of PT activation and function remain incompletely understood. It is very likely that physically PT can occur through several pathways (Bonora *et al*., 2022; Bround *et al*., 2020). One of the key challenges in the field is understanding the roles of the ATP synthase and ANT in this process. Compelling evidence from several independent laboratories supports competing interpretations suggesting that a core part of PTP involves either the ATP synthase complex or ANT, both of which could be transformed into the high-conductance pore (Alavian *et al*., 2014; Brustovetsky and Klingenberg, 1996; Giorgio *et al*., 2013). In both cell types lacking either ATPase or ANT, calcium treatment causes calcium release and membrane depolarization that is inhibited by CSA (Carroll *et al*., 2019; Karch *et al*., 2019). Here, using the same knock-out cell models, we observed the phenomena of membrane depolarization. However, holographic imaging revealed that mitochondria in these mutant cells did not undergo high amplitude permeabilization. This suggests that both ATP synthase and ANT are essential for the development of the high-conductance PTP. These results are in agreement with the previously proposed model that, in fact, the functional PTP complex would require presence of both ATPase and ANT (Halestrap et al., 2002). The fact that none of these complexes are required for Ca-induced mitochondrial depolarization would explain the controversy in the literature regarding their roles in PTP.

In summary, the two phenomena observed in our experiments suggest the presence of a low-conductance mode of PTP, which occurs independent of the ATP synthase and ANT, which are required for the high-conductance mode of PTP. We hypothesize that PTP development might be a two-channel phenomenon (ANT and ATPase) that demonstrate interdependence. The lack of PTP in cells lacking ATPase and ANT opens an exciting possibility that the two steps of PTP might involve different molecular structures. It is tantalizing to suggest that selectively targeting ATP synthase and ANT might help to identify compounds that would prevent mitochondrial high amplitude permeabilization but allow for a protective depolarization step, which would prevent mitochondria from toxic calcium overload and oxidative stress.

## Acknowledgements

We thank Prof. Mike Murphy (University of Cambridge, UK) and Prof. Nickolay Brustovetsky (University of Indiana, USA) for critical reading of the manuscript and fruitful discussion and Prof. John Walker (University of Cambridge, UK) for providing HAP 1 cell lines and Prof. Jeff Molkentin for providing MEF cell lines. This work was supported by grants # R35GM139615 and R01GM115570 from NIGMS (to E.P.) and AHA Postdoctoral fellowship (to M.N.).

