## Supplementary material for "Both ANT and ATPase are essential for mitochondrial permeability transition but not depolarization": Fig. S

### Slide 1
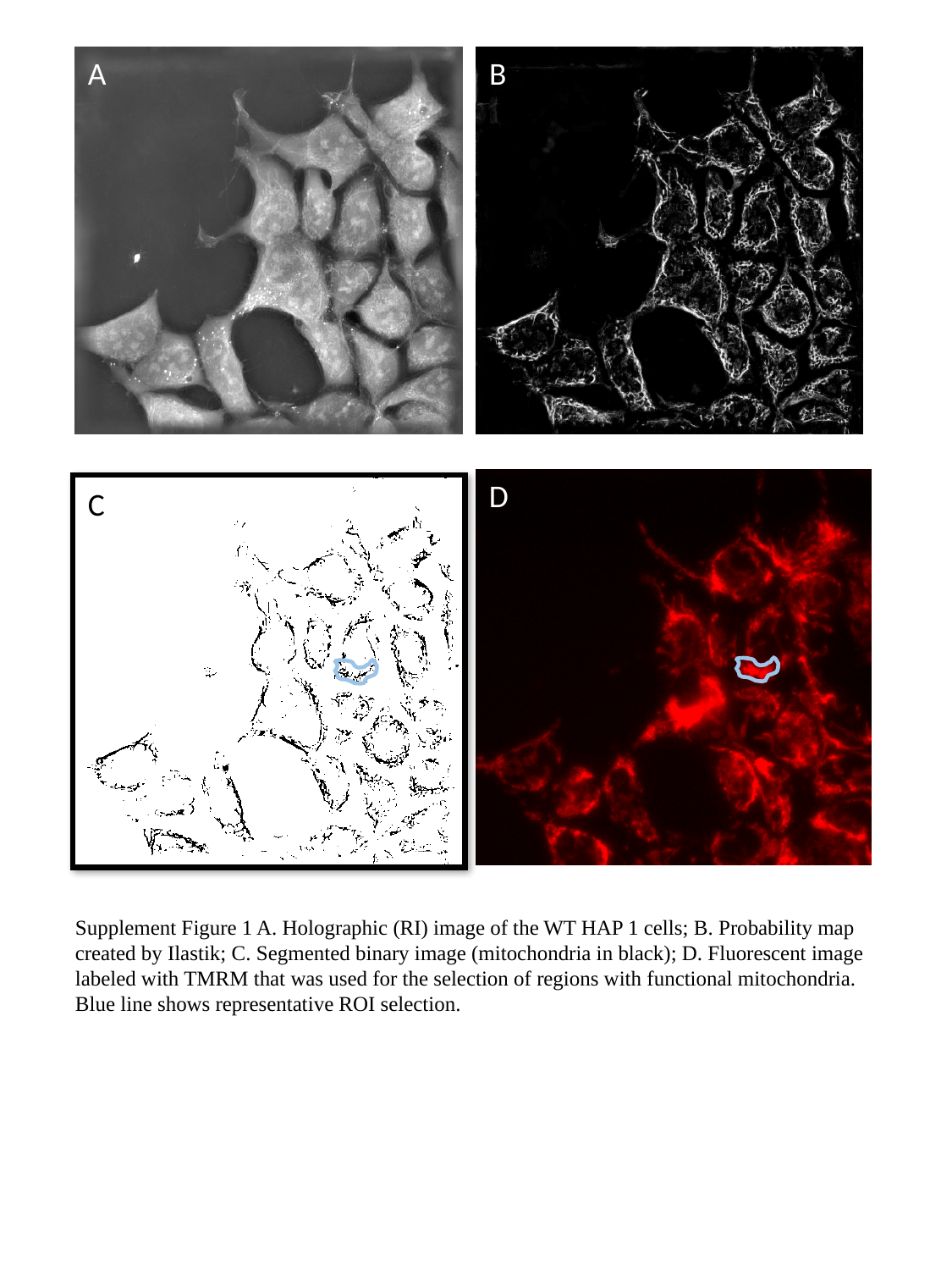

A
B
D
C
Supplement Figure 1 A. Holographic (RI) image of the WT HAP 1 cells; B. Probability map created by Ilastik; C. Segmented binary image (mitochondria in black); D. Fluorescent image labeled with TMRM that was used for the selection of regions with functional mitochondria.
Blue line shows representative ROI selection.

### Slide 2
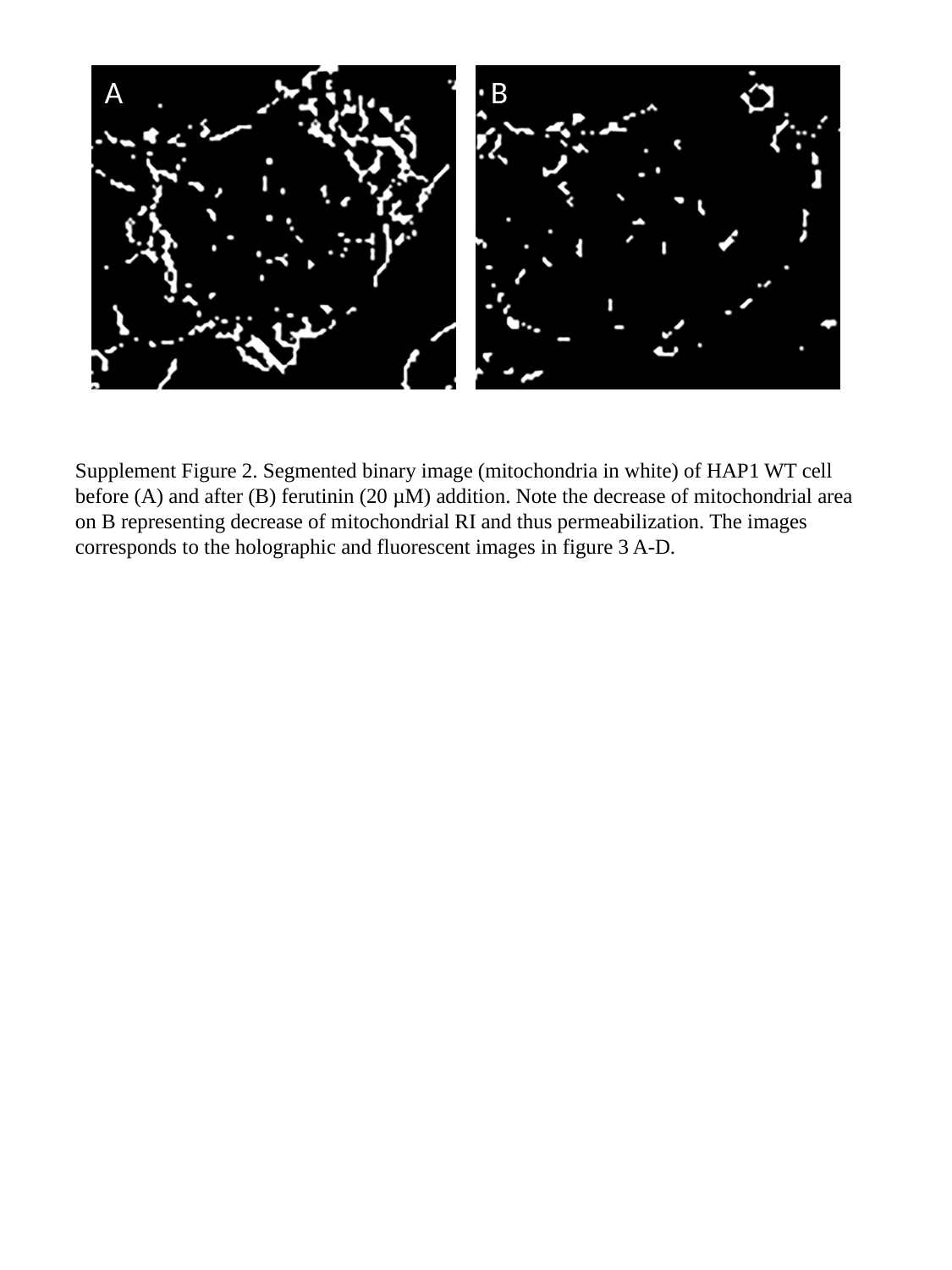

A
B
Supplement Figure 2. Segmented binary image (mitochondria in white) of HAP1 WT cell before (A) and after (B) ferutinin (20 µM) addition. Note the decrease of mitochondrial area on B representing decrease of mitochondrial RI and thus permeabilization. The images corresponds to the holographic and fluorescent images in figure 3 A-D.

### Slide 3
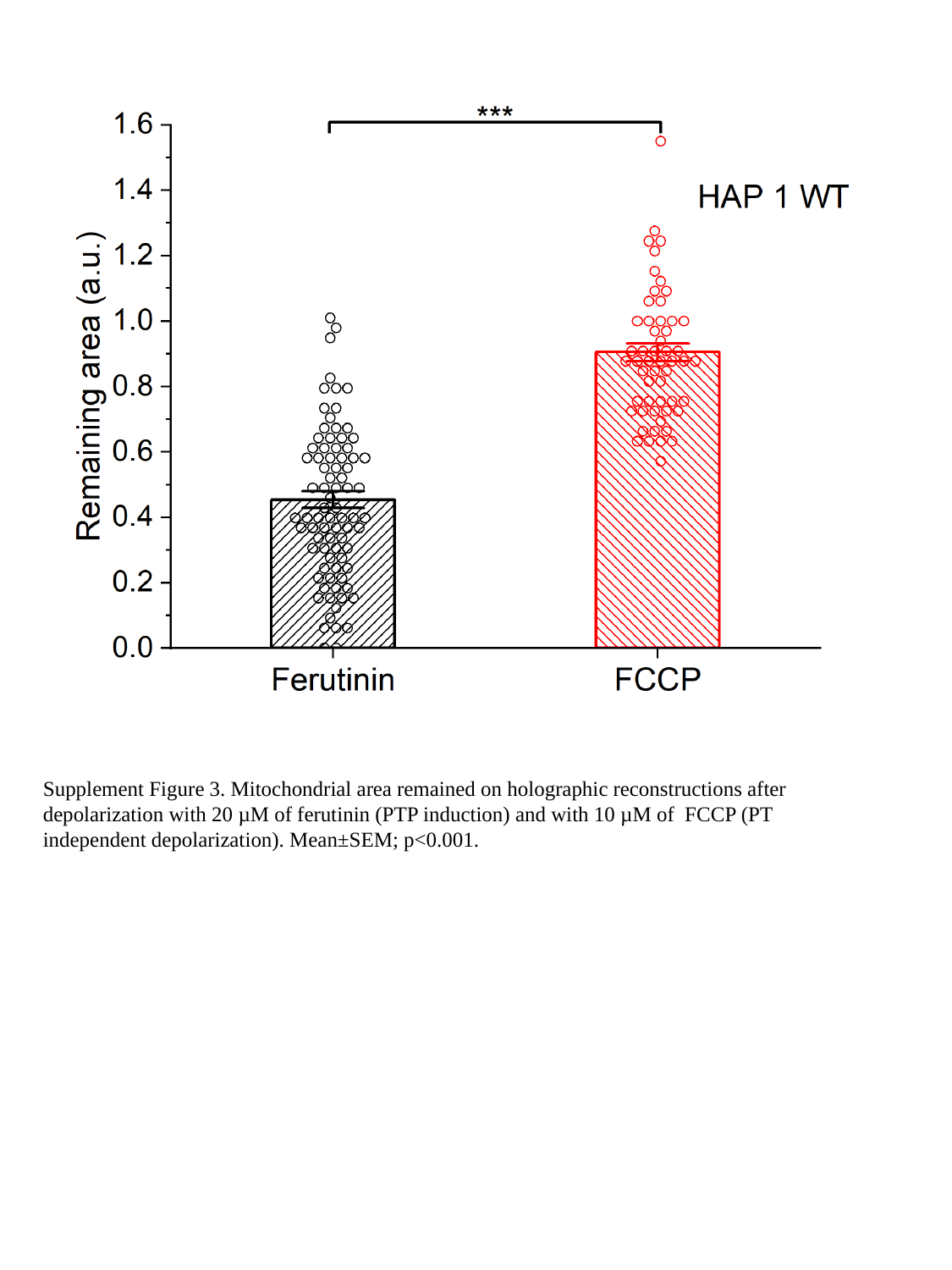

Supplement Figure 3. Mitochondrial area remained on holographic reconstructions after depolarization with 20 µM of ferutinin (PTP induction) and with 10 µM of FCCP (PT independent depolarization). Mean±SEM; p<0.001.

### Slide 4
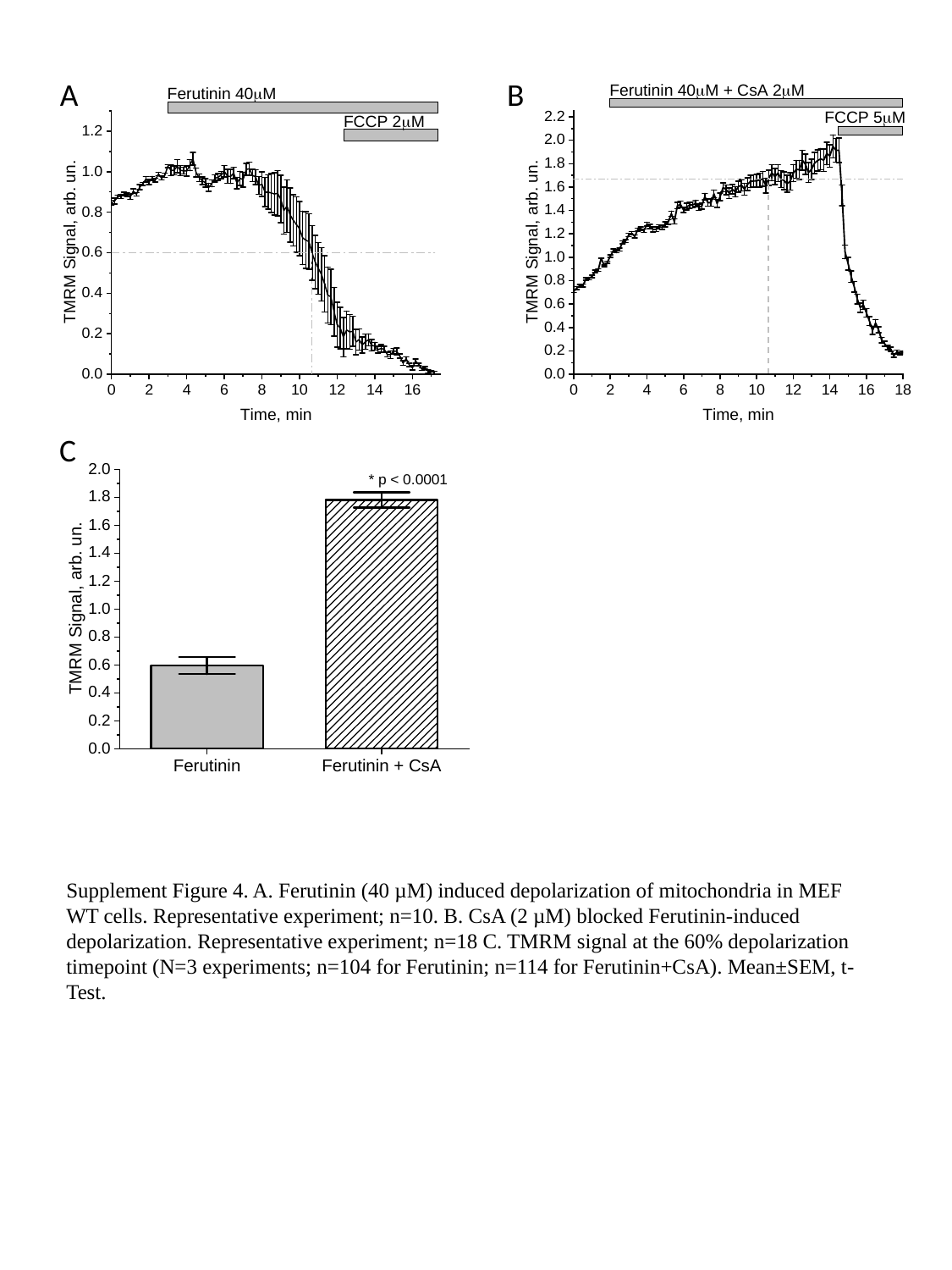

A
B
C
Supplement Figure 4. A. Ferutinin (40 µM) induced depolarization of mitochondria in MEF WT cells. Representative experiment; n=10. B. CsA (2 µM) blocked Ferutinin-induced depolarization. Representative experiment; n=18 C. TMRM signal at the 60% depolarization timepoint (N=3 experiments; n=104 for Ferutinin; n=114 for Ferutinin+CsA). Mean±SEM, t-Test.
